# Cortical Temporal Integration Window for Binaural Cues accounts for Sluggish Auditory Spatial Perception

**DOI:** 10.1101/2021.12.14.472656

**Authors:** Ravinderjit Singh, Hari M. Bharadwaj

## Abstract

The auditory system has exquisite temporal coding in the periphery which is transformed into a rate-based code in central auditory structures like auditory cortex. However, the cortex is still able to synchronize, albeit at lower modulation rates, to acoustic fluctuations. The perceptual significance of this cortical synchronization is unknown. We estimated physiological synchronization limits of cortex (in humans with electroencephalography) and brainstem neurons (in chinchillas) to dynamic binaural cues using a novel system-identification technique, along with parallel perceptual measurements. We find that cortex can synchronize to dynamic binaural cues up to approximately 10 Hz, which aligns well with our measured limits of perceiving dynamic spatial information and utilizing dynamic binaural cues for spatial unmasking, i.e. measures of binaural sluggishness. We also find the tracking limit for frequency modulation (FM) is similar to the limit for spatial tracking, demonstrating that this sluggish tracking is a more general perceptual limit that can be accounted for by cortical temporal integration limits.

## 1 Introduction

Temporal information is encoded with microsecond precision in the auditory periphery and is progressively transformed into a rate-based code (Joris et al., 2004). However, cortical neurons can also synchronize to input auditory cues, albeit at lower frequencies than subcortical structures (Liang et al., 2002; Joris et al., 2004; Bendor and Wang, 2008). The impact of this cortical synchronization on perception is poorly understood. Studies have found cortical synchronization to be useful in explaining perception of time-reversed animal vocalizations, encoding of stimulus onset, representation of auditory objects, and the coding of amplitude modulations (AMs) (Walker et al., 2008; Schnupp et al., 2006; Phillips et al., 2002; Heil, 2004; Elhilali et al., 2009; Bendor and Wang, 2007). As the modulation rates (frequencies) of sounds are increased, the elicited percept changes from that of being able to perceptually follow discrete events (i.e., the individual peaks and troughs of the modulation), to a flutter, and then to a pitch. Indeed, correlates of these qualitative changes in the percept of AM can be found in the temporal synchronization capabilities of different cortical regions (Liang et al., 2002; Bendor and Wang, 2007, 2008). We demonstrate here that analogous qualitative perceptual switches also occur in the processing of frequency modulation (FM) and binaural modulation (BM; fluctuations in binaural cues). The perceptual switch seen with BM processing has been called “binaural sluggishness” in the literature, because it was thought that this sluggishness phenomenon may be unique to the binaural system. Instead, here, we explore the possibility that cortical synchronization capabilities may place common neurophysiological constraints leading to sluggish processing of a range of auditory cues, including monaural AM and FM (Fig. 1). Our experiments demonstrate that the binaural temporal analysis/integration window of the neural population in a hierarchically later cortical region (measurable with a latency of about 100 ms with electroencephalography; EEG), can quantitatively account for the sluggish perception seen in BM processing.

**Figure 1.**
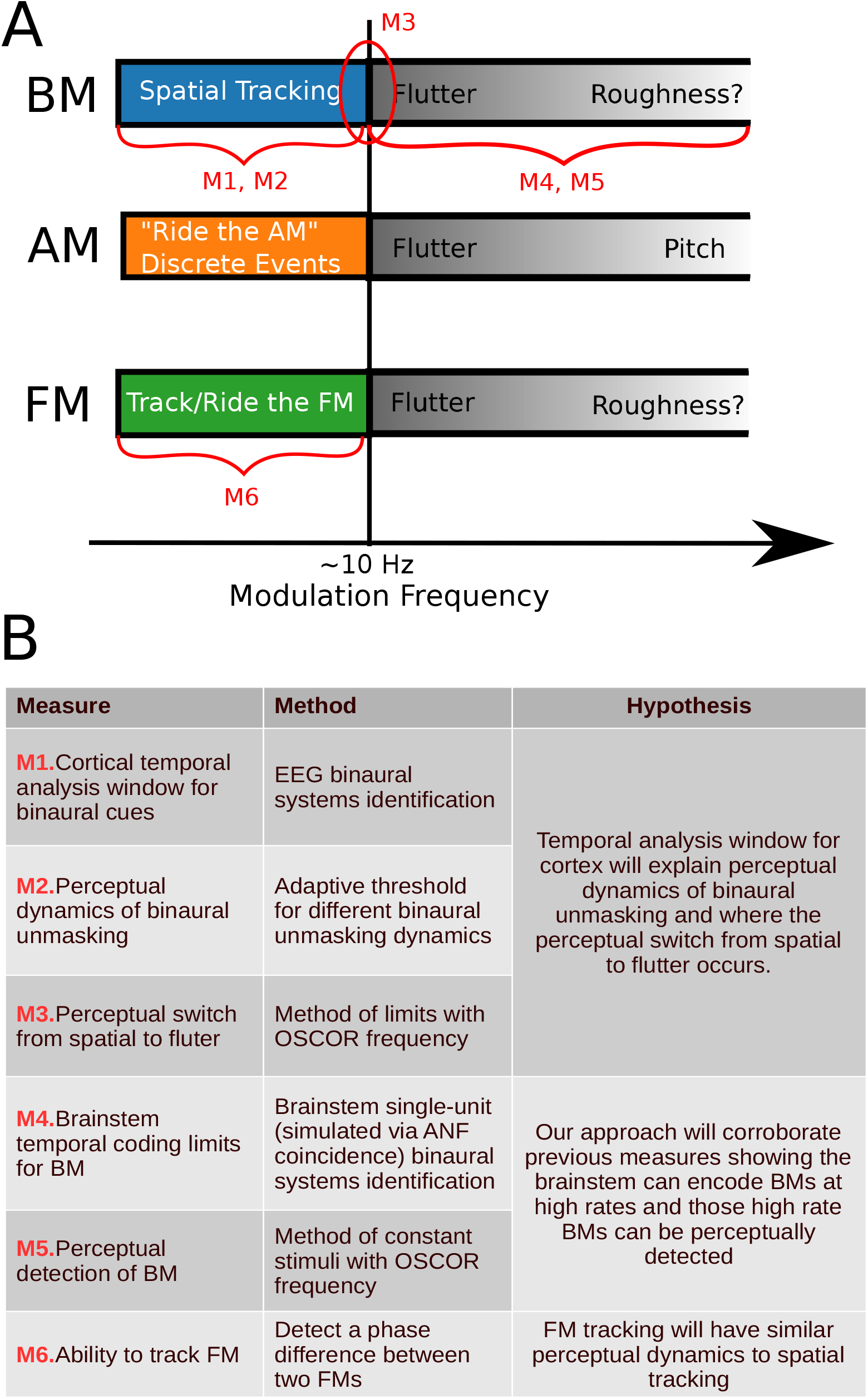
**A**. This figure depicts the similarity in temporal switches in perception that occur between binaural modulation (BM), amplitude modulation (AM), and frequency modulation (FM). In red, the measures taken in this work are depicted and laid out in the table shown in **B**.

In this investigation, BMs are applied to two binaural cues based on the temporal fine structure in sounds, namely interaural time delay (ITD) and interaural correlation (IAC). ITD is the difference between the arrival times of corresponding components of a sound in the two ears, and is useful for sound lateralizarion; IAC is the correlation between sounds reaching the two ears, and can directly influence the perceived spatial extent/width of a sound. Both ITD and IAC can be useful cues for listening in noisy environments (Hawley et al., 2004; Lingner et al., 2016; Robinson and Jeffress, 1963). How binaural cues, e.g. ITD and IAC, are processed when they are dynamic, is currently not well understood. Single-unit data from the brainstem shows that cells can encode BMs in ITD and IAC in the 100s of Hz (Joris et al., 2006; Siveke et al., 2008; Zuk and Delgutte, 2017; Joris, 2019). Single-unit data from primary auditory cortex (A1) shows neurons can synchronize to binaural beats, a dynamic interaural phase difference (IPD), up to a median synchronization rate of about 20 Hz (Fitzpatrick et al., 2009; Scott et al., 2009). Behaviorally, humans can detect BMs in the 100s of Hz (Grantham, 1982; Grantham and Wightman, 1978; Siveke et al., 2008); however, studies probing the use of binaural cues to perform a spatial task, e.g. spatial unmasking, have found that humans only benefit from low-rate BMs, below 10 Hz (Culling and Colburn, 2000; Culling and Summerfield, 1998; Grantham and Wightman, 1979; Kollmeier and Gilkey, 1990; Shackleton and Bowsher, 1989). These binaural unmasking studies led to the notion of “binaural sluggishness” in the literature as the binaural system seemed particular slow in comparison to human ability to detect monaural cues. However, a broader view of the literature suggests a dichotomy where tasks that rely on the percept being spatialized in nature appear slow, while simple detection of binaural fluctuations can be more than an order of magnitude faster. Indeed, dynamic spatial percepts have been anecdotally reported to become a flutter at frequencies above approximately 7-10 Hz (Grantham and Wightman, 1978; Siveke et al., 2008; Zuk and Delgutte, 2017). Thus, BMs seem to have analogous temporal perceptual limits as AMs, in that both demonstrate a qualitative switch around 7-10 Hz, Fig. 1; with BMs, the percept switches from spatialized to a mere flutter, and with AMs, the percept switches from being able to “ride” the AM (or perceive individual peaks and troughs discretely), to also perceiving a flutter. This dichotomy (fast and slow) in the behavior is accompanied by evidence of fast temporal processing in the brainstem and slower temporal processing in the cortex. However, the slower temporal processing found in cortex thus far, particularly primary auditory cortex (median synchronization limits of 20 Hz), is not as slow as spatial perception observed in human behavior (Fitzpatrick et al., 2009; Scott et al., 2009); however, in this comparison, it should be acknowledged that human perceptual limits are being compared to non-human physiology.

We evaluate how cortical binaural synchronization in humans explains human dynamic binaural perception. A temporal analysis window is quantified using EEG with a novel binaural systems identification technique which modulates IAC or ITD with a maximum length sequence (m-seq). Using the same system identification technique, we also estimate brainstem binaural synchronization limits in chinchillas to both corroborate previous brainstem measurements and to validate our novel approach against prior single-unit data obtained using conventional approaches. We conduct three behavioral experiments (1) to characterize detection limits for BMs, (2) estimate the frequency limits at which the perceptual switch from dynamic spatial to a non-spatial flutter occurs, and finally (3) to quantify the perceptual dynamics of binaural unmasking; we then compare the behavioral frequency functions to synchronization limits of cortex and brainstem. To complement our binaural measures, we also measured subjects’ ability to monaurally detect a phase difference between two spectrally distant FMs, which requires being able to temporally follow the individual cycles of the FMs to test whether the ability to temporally track BMs (i.e. spatial tracking) ceases at similar modulation rates as for other auditory cues. The different experiments conducted, and hypotheses tested, are outlined in Fig. 1. Our results reveal a neural source with a latency of ~100 ms that synchronizes to BMs slower than 10 Hz, and can quantitatively account for the sluggish dynamics seen in spatial unmasking. We also find that BM and FM stimuli can be perceptually tracked out to similar modulation frequencies (approximately 10 Hz). These results suggest the temporal response properties of later (~100 ms latency) areas of cortex may constrain our ability to perceptually track various dynamic auditory cues.

## 2 Results & Discussion

In the present study, we employed a novel approach to characterize neural coding of binaural modulations. Neuroscience in general, and auditory neuroscience in particular has a rich history of characterizing temporal coding. Indeed, temporal modulation transfer functions (tMTFs) have been measured from various levels of the auditory system for spectral, amplitude, and binaural modulations (Joris et al., 2006; Liang et al., 2002; Zuk and Delgutte, 2017). The conventional approach involves playing sinusoidal modulations (i.e., a single frequency in the modulation domain), and measuring neural phase locking as a function of the modulation frequency. This approach is time consuming, as many frequencies need to be measured individually. Moreover, given the highly non-linear and adaptive nature of the central auditory neural response, the results from a discrete single-frequency sampling approach may miss interesting characteristics that are unique to the broadband nature of real-word stimuli. To mirror the complex broadband modulation profile encountered in the environment, we applied a broadband binaural modulation using a modified maximum length sequence or m-sequence (m-seq). Throughout, we refer to this modified stimulus as the “extended m-seq” (em-seq) (Fig. 4). This approach allows us to simultaneous measure the coding across all modulation frequencies of interest with one ongoing stimulus, and provides a significant improvement in experimental efficiency compared to traditional tMTF measurements.

Our measurement using a extended m-seq (em-seq) and subsequent PCA analysis to obtain a “source binaural temporal response function” (sBTRF) is explained in detail in the methods section. The sBTRF for IAC and ITD is shown in Fig. 2. The sBTRF reflects how the dominant underlying cortical sources responds to dynamic binaural stimuli. The PCA weights are depicted in the topomaps in Fig. 2 which suggests bilateral sources in the auditory cortex. Neuroimaging data in humans and data from animals support the notion that perceived auditory space is computed in auditory regions in the temporal lobe, but beyond primary auditory cortex (Lewald and Getzmann, 2011; Viceic et al., 2006; van der Zwaag et al., 2011; Stecker et al., 2005). Consistent with this, the dominant group delay estimated from the sBTRF was approximately 100 ms for both IAC and ITD. Studies on the encoding of dynamic binaural cues in cortex have mainly focused on neurons in the primary auditory cortex and have reported median synchronization limits of approximately 20 Hz (Fitzpatrick et al., 2009; Scott et al., 2009); the 20 Hz limit is much faster than what is observed in human behavior. The magnitude response of the sBTRF for IAC loses 6 dB or half its amplitude by 5 Hz and the mean response falls into the noise floor at approximately 9 Hz, while for ITD, the amplitude peaks around 4.5 Hz and falls into the noise floor at approximately 9 Hz as well. This slower frequency limit estimated from the sBTRF further corroborates the notion that the dominant contributors may be sources that are hierarchically downstream to the primary auditory cortex.

**Figure 2.**
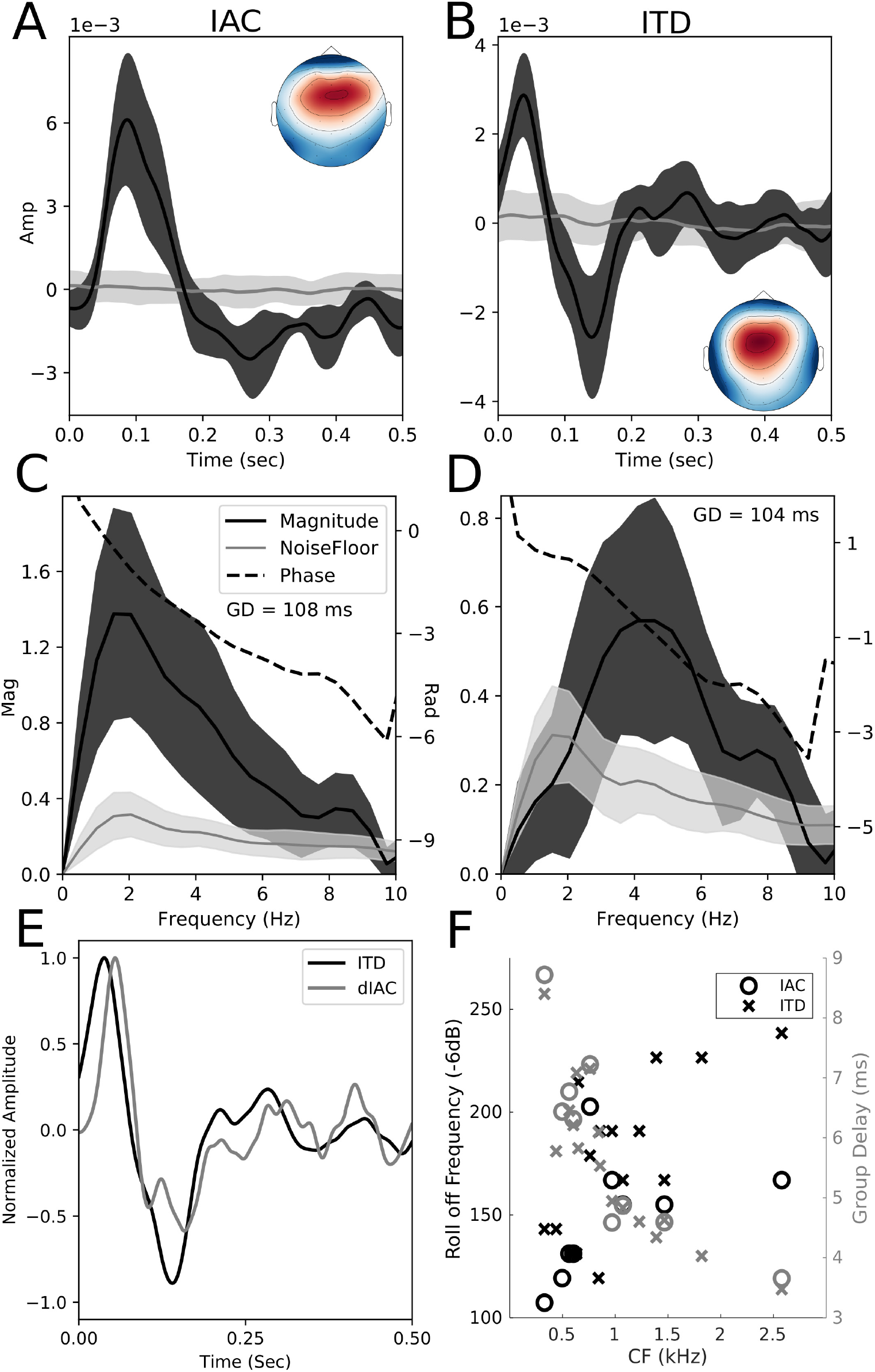
In **A-D**, the solid line is the mean response and shading represents the 95% confidence interval calculated using the standard error. **A-B** shows the source binaural temporal response function (sBTRF) for IAC and ITD as well as the topomap for the sBTRF. **C-D** shows the frequency response of the sBTRF for IAC and ITD. The data was high-passed at 1 Hz. **E** shows a plot of the ITD sBTRF as well as the derivative of the IAC sBTRF with the amplitudes normalized between 1 and −1. **F** shows the roll off frequency and group delay for the brainstem responses simulated from ANF coincidence.

The sBTRF shapes for IAC and ITD have an interesting relationship in that the shape of the ITD sBTRF roughly matches the derivative of the IAC sBTRF (Fig. 2D). This relationship can be explained in the context of a two-channel population code model for sound location(Stecker et al., 2005; Salminen et al., 2009; Magezi and Krumbholz, 2010). With IAC, it is likely all cortical binaural cells (in both channels) respond to IAC the same way, exhibiting a smaller response at a low IAC and a larger response at high IAC. In contrast, with ITD, two-channel models considered in the literature posit that one channel would respond more favorably to sound locations on one lateral side vs the other (Stecker et al., 2005; Salminen et al., 2009; Magezi and Krumbholz, 2010). Therefore with the ITD em-mseq (which bounced between two azimuths), one channel may be much more active for one azimuth and the other more active for the other azimuth leading to a derivative-like response compared to the response seen for IAC. Fig. 2F show results from single-unit measurements in chinchillas, demonstrating that the approach utilizing an em-mseq can also be readily adapted to spiking data. We estimated binaural brainstem responses from a coincidence analysis on measurements from the auditory nerve recordings in chinchillas. Previous studies have shown that brainstem response properties can be reliably predicted from nerve responses using this approach (Joris et al., 2006). A drop of 6dB in power was used as the synchronization limit to be consistent with previous literature (Joris et al., 2006; Zuk and Delgutte, 2017). Our brainstem estimates indicate synchronization up to 100s of Hz, in line with previous measurements (Joris et al., 2006; Siveke et al., 2008; Zuk and Delgutte, 2017).

Results from behavioral experiments are shown in Fig. 3. Detection thresholds for BMs were measured using a binaural oscillating-correlation (OSCOR) stimulus, where the IAC was varied sinusoidally at different BM rates. Results revealed that humans can detect BMs as fast as 100s of Hz, consistent with our ability to temporally encode fast BMs in the brainstem. Importantly, being able to *detect* BMs does not mean that the listener is making use of fluctuations in perceived lateralization for all rates; just that changing IAC could be discriminated from fixed. It should be noted that experiments were primarily conducted with noise stimuli that were bandlimited between 0.2-1.5 kHz to match the range of frequencies over which humans have high sensitivity to ITDs (Brughera et al., 2013), perhaps constrained by the limits of neural phase locking (Verschooten et al., 2019). We also measured OSCOR BM detection in one participant with white noise and observed that detection extended to higher BM rates (S.3). To quantify the anecdotally reported phenomenon that the perception of BMs switches from being spatial to a flutter between 6-10 Hz (Grantham and Wightman, 1978; Siveke et al., 2008; Zuk and Delgutte, 2017). We measured this in one participant and found the switch from spatial to flutter occurred at 9.3 Hz in that participant.

**Figure 3.**
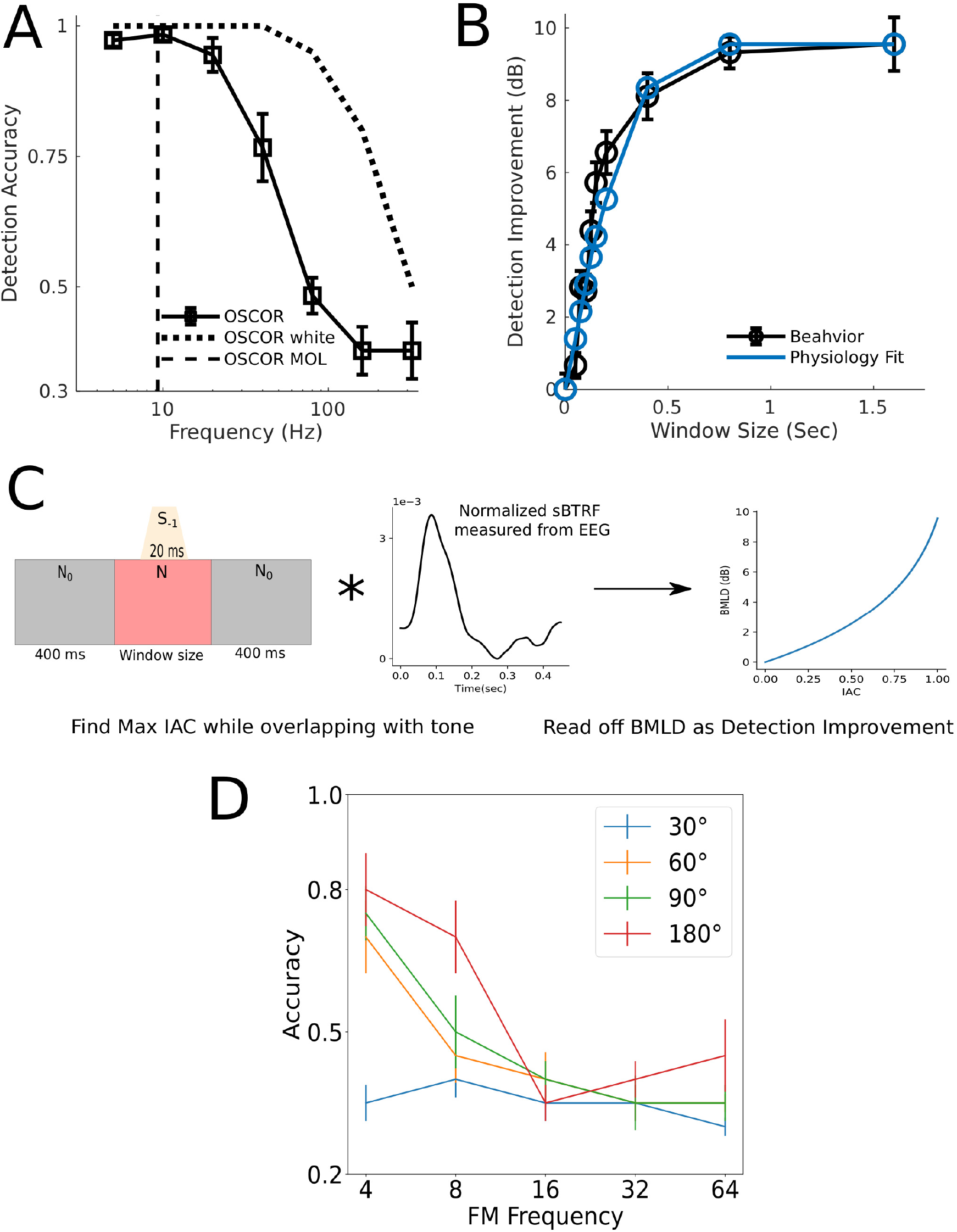
**A** shows results from all experiments using the OSCOR stimulus. **B** shows the results for a binaural unmasking paradigm as well as how well the IAC sBTRF window fits the behavioral data. **C** demonstrates how the physiology fit was obtained and is based on procedures from Culling and Summerfield (1998). **D** shows the results from our FM phase difference detection experiment.

In contrast to detection experiments, binaural unmasking experiments rely on subjects’ ability to use brief periods of perceived spatial separation (i.e., differences in lateralization/location) between target and masking sounds to derive masking release (Culling and Summerfield, 1998). Indeed, the results of our experiment measuring the dynamics of binaural unmasking reveals the sluggish nature of spatial processing; the binaural masking release (measured as dB improvement in target detection) continues to grow even as the duration of the spatial separation is increased in the range of tenths of a second, Fig. 3 B,C. Strikingly, when we used the physiologically measured sBTRF for IAC as the temporal integration window, the predicted masking release shows excellent correspondence to the behavioral data, with the mean-squared error (MSE) of the fit being as small as 0.6 dB^2^ (this low MSE is despite excluding the end points, i.e., 0 and the asymptotic value at 1.6 seconds which were constrained to match the measurements). Therefore, the sluggishness in binaural perception is well explained by the physiological properties of cortical processing, specifically that of a cortical source that is measurable using EEG with a group delay of approximately 100 ms. Data from primary auditory cortex has shown median synchronization limits to binaural cues to be 20 Hz (Fitzpatrick et al., 2009; Scott et al., 2009) which is faster than our measure here that fits the behavior, suggesting this sluggishness arises in a hierarchically later cortical source (consistent with the longer 100 ms latency) and not primary auditory cortex. Finally, to test whether a monaural percept can also exhibits sluggishness, we measured the ability to detect a phase difference between FMs at the same rate imposed on spectrally distant carriers. This task was chosen because it relies on subjects’ ability to perceive discrete variations in the modulation and temporally track it (i.e., being able to “ride” the peaks and troughs) to detect the phase difference. This is analogous to being able to track the changing lateratization or spatial location with BM stimuli. We hypothesized that similar to BMs, FMs would show the same tracking limits. Indeed, we observed that FMs can be temporally tracked up to about 10 Hz, similar to our BM results, Fig. 3 D.

In summary, the sBTRF measured for IAC and ITD provides a neural correlate for the sluggish perception of auditory space. Furthermore, the latency of the dominant source contributing to the sBTRFs suggests the perceptual limits arise from auditory cortical regions downstream of the primary auditory cortex. Previous investigations focusing on the brainstem and midbrain processing have not produced any candidate correlates for this spatial sluggishness (Joris et al., 2006; Siveke et al., 2008; Shackleton and Palmer, 2010; Zuk and Delgutte, 2017, 2019; Joris, 2019). Cortical single-unit measurement in A1 have revealed lower limits of temporal coding than the brainstem, but not low enough to account for the sluggish the behavioral estimates of sluggish spatial perception in humans (Fitzpatrick et al., 2009; Scott et al., 2009). The shape of the sBTRF measured here yielded slugggishness estimates that were a very close match to the spatial sluggishness seen in binaural unmasking. Our results also make it clear that the processing of binaural cues is not uniquely sluggish, and that monaural temporal perception also exhibits sluggishness when the behavioral tasks are set up to probe analogous aspects of temporal tracking. This suggests that “binaural sluggishness” may be somewhat of a misnomer. Indeed, with AM, fluctuations can be followed only up to about 10 Hz if individual acoustic events are to be discretely perceived; then at higher rates, an acoustic flutter is heard (similar to the binaural flutter), and then finally a pitch (Bendor and Wang, 2007). Correlates of these qualitative changes in AM perception can be found in the temporal coding properties of different regions along the auditory pathway (Bendor and Wang, 2007, 2008). Here, we demonstrated that a similar 10 Hz limit also exists in the tracking of FMs. Indeed, given that hierarchical cortical processing occurs via highly conserved columnar circuits (Felleman and Van Essen, 1991), and that cortical neurons generally exhibit tuning to a range of different acoustic cues at the single-unit level, it is reasonable to expect that the temporal coding properties of cortical neurons would constrain the processing of many different cues similarly. In this view, the sluggishness in tracking a moving sound (spatial sluggishness) would be no different than the sluggishness in tracking discrete amplitude or frequency fluctuations. Rather, cortical processing limits may impose a general sluggishness in the processing of a range of auditory cues. Our results also suggest that although the temporal integration window progressively expands as we ascend the auditory pathway (progressive temporal to rate transformation), the temporal synchronization properties of higher stages of cortical processing can still directly relate to certain aspects of perception. The perceptual significance of this low frequency temporal processing is under investigated, and could be important for our understanding of the mechanisms supporting the perception of complex acoustic scenes. Our em-seq approach to evaluate how well cortex can track a particular auditory cue can be readily extended to other acoustic cues, such as AM, to investigate cortical temporal coding more broadly. Finally, to the extent that temporal response properties of higher-order areas relate to perception of dynamic scenes, individual differences in physiology measured using the em-seq approach may also be useful for predicting individual listening outcomes in complex everyday environments.

## 3 Methods & Materials

We conducted six experiments, illustrated in Fig. 1B, to investigate how temporal processing in cortex relates to binaural perception and to evaluate whether a common sluggishness phenomenon can be found across the processing of many auditory cues. Two experiments involved electrophysiological measurements using our novel binaural systems identification approach to quantify the temporal coding properties of brainstem and cortex for binaural modulations. Four experiments were behavioral studies in humans designed to probe the temporal limits of different binaural perceptual abilities, and to test whether physiologically measured limits can predict perception.

### 3.1 Human Participants

Nine participants (2 female) with an average age of 25 (18-34) were recruited from the greater Lafayette area through posted flyers and advertisements. Audiograms were measured using calibrated Sennheiser HDA 300 headphones by employing a modified Hughson-Westlake procedure. All subjects had hearing thresholds better than 25 dB HL in both ears at octave frequencies between 250 – 8 kHz. All subjects provided informed consent, and all measurements were made in accordance with protocols approved by the Internal Review Board, and the Human Research Protection Program at Purdue University.

### 3.2 EEG Recording

Digital stimuli were designed with custom scripts in Matlab (The MathWorks Inc., Natick, MA) at a sampling rate of 48828.125 Hz, and converted to analog voltage signals using an RZ6 audio processor (Tucker-Davis Technologies, Alachua, Florida). The voltage signals were converted to sounds and delivered to the ears via ER2 insert earphones (Etymotic Research, Elk Grove Village, IL) coupled to foam ear tips. EEG measurements were made with a 32-channel system (Biosemi Active II system, Biosemi, Amsterdam, Netherlands). EEG data was sampled at 4096 Hz and filtered between 1-40 Hz, and then downsampled to 2048 Hz. EEG data were re-referenced to the average of all electrodes. Ocular artifacts were removed using the signal-space projection approach (Uusitalo and Ilmoniemi, 1997). Projections were manually applied for each participant based on the topographic pattern of the noise-space weights which were manually chosen to correspond to the expected pattern of blink and saccade artifacts. EEG data were collected as stimuli were played passively with participants watching a muted video of their choosing with subtitles. To remove movement artifacts, trials that had peak to peak deflections greater than 200 *μ*V were then rejected.

### 3.3 Auditory Nerve Responses to Simulate Binaural Processing

Auditory nerve responses were collected in chinchillas. Male chinchillas (N=5) weighing 400 – 650 grams were used in accordance with protocols approved by the Purdue Animal Care and Use Committee. Binaural processing was simulated by playing the stimulus intended for each ear separately and recording from the same nerve fiber, and then performing a coincidence analysis. Coincident spikes (within 50 *μ*s bins) across the “left” and “right” spike train was treated as amounting to a binaural spike. Previous work has shown that this coincidence analysis approach can reliably estimate the response properties of binaural cells in the midbrain (Joris et al., 2006). All procedures laid out in this section were approved by the Purdue University Animal Care and Use Committee.

Anesthesia was induced with xylazine (1-2 mg/kg s.c.) and ketamine (60-65 mg/kg, s.c.). Anesthesia was maintained with ketamine (20-40 mg/kg, i.m.) and diazepam (1-2 mg/kg, i.m.) through intramuscular injections every 2 hours as indicated by the presence of reflexes for the duration of the experiment (10-16 hours). A heating blanket and rectal thermometer were used to regulate and monitor body temperature throughout the experiment. The skin and muscles overlying the skull were reflected to expose the bony ear canals and bullae. Hollow ear bars were placed close to the tympanic membrane. The AN bundle was exposed at its exit from the internal acoustic meatus via a posterior-fossa craniotomy and aspiration cerebellotomy. Acoustic stimuli were presented monaurally through an ear bar using Etymotic ER2 and calibrated using a probe microphone placed within a few mm of the tympanic membrane (Bruel and Kjaer 4182). Glass pipettes with impedance ranging between 60-90 MΩ were advanced into the auditory nerve using a hydraulic microdrive. Recordings were amplified, band-pass-filtered and stored on a PC. Spikes were identified using a time-amplitude window discriminator, and spike times were stored with 10 *μ*s resolution.

Single fibers were isolated by monitoring the spike recording via an oscilloscope and listening for spikes while noise pips were played. After isolating a fiber, its tuning curve was measured using an automated procedure that played a series of tone pips and measured the minimum level required to evoke 1 more spike than a subsequent 50 ms silent period (Chintanpalli and Heinz, 2007). Units where we were unable to measure at least 20 repetitions of the left and right stimulus were excluded from analysis.

### 3.4 Novel Binaural Systems Identification using Maximum Length Sequences

Systems identification approaches are widely used in Neuroscience (Eggermont, 1993; Ringach and Shapley, 2004). In audition, systems identification approaches are useful for determining the frequency responses of a system, both at single-neuron and population levels (Henry et al., 2019; Recio-Spinoso et al., 2005). Binaural systems identification has been used to study the temporal coding limits of the binaural system primarily through temporal modulation transfer functions (tMTF) computed from calculating phase locking to individual frequencies of binaural modulation (Fitzpatrick et al., 2009; Joris et al., 2006; Scott et al., 2009; Siveke et al., 2008; Zuk and Delgutte, 2017). However, the approach of constructing a tMTF from responses to individual frequencies is experimentally inefficient. Also central neural responses can be sensitive to the input statistics of a stimulus, so an approach that better mimics the broadband nature of real-world stimuli may capture interesting characteristics that a discrete single-frequency sampling approach may miss (Wang, 2007; Theunissen et al., 2000). Here we use a novel binaural systems identification technique where we modulate a binaural cue using a maximum length sequence (m-seq) allowing us to evaluate a continuous range of frequencies using a single stimulus.

Maximum length sequences (m-seqs) have been used to map the receptive fields of visual neurons, identify the cues used in visual tracking, obtain auditory brainstem responses, and characterize room acoustics (Aptekar et al., 2014; Chu, 1990; Reid et al., 1997; Shi and Hecox, 1991; Burkard et al., 2017). An m-seq is a signal that only takes one of two discrete values (e.g., +1 or −1). It is a pseudorandom sequence constructed through a series of feedback shift registers of *n* bits, giving the full sequence a length of 2^*n*^ − 1. An m-seq has a flat spectrum, similar to white noise, but has a lower crest factor in that its values are bounded on both ends, making it attractive for systems identification. By using the m-seq as the input to a system and then cross-correlating it with the output of the system, an estimate of the impulse response can be obtained. For more details on the m-seq, we suggest the following sources (Aptekar et al., 2014; Chu, 1990; Reid et al., 1997; Shi and Hecox, 1991).

The m-seq has a nearly white spectrum; however auditory responses to binaural modulation, especially in cortex are expected to be slow. Indeed, previously measures from the primary auditory cortex of rabbits and macaques suggest that responses roll off around 20 Hz (Fitzpatrick et al., 2009; Scott et al., 2009). Thus, if we played the m-seq as it is conventionally constructed, its energy would be spread out up to *fs/*2, which is over 20 kHz in this case, because our sampling rate (*fs*) is 48828.125 Hz. This is orders of magnitude wider that than expected frequency response range of the system resulting in much of the stimulus energy for system characterization being spent without eliciting a measurable response component, leaving only a small fraction to characterize the system in the region that it is active (e.g., below 20 Hz). Therefore we modified the m-seq to an extended m-seq (em-seq) to have a sinc-function-shaped spectrum instead of a white spectrum, so that most of the characterization energy of the m-seq is in the region of interest for the system being characterized. The sinc-function shape will modulate the measured system response; however, the em-seq can be designed so that it’s frequency response is mostly flat in the region the underlying system is active. The procedure to construct the em-mseq is explained next.

The em-mseq is constructed by elongating the duration of each point in the conventional m-seq to create an extended m-seq, i.e., em-mseq. In the conventional construction, each point in the m-seq is 1*/fs* in duration where fs is the sampling rate. By elongating that duration of that state to T, the em-seq spectrum instead of being flat, takes a sinc-function shape, with only minimal energy above *f* = 1*/T*, but approximately flat (losing less than 2 dB) between *f* = 0, and *f* = 1/2*T*. For example, if characterizing a system that is expected to be active up to 40 Hz, elongating each point to a duration of 12.5 ms would be appropriate because then the resulting sinc spectrum will be fairly flat up to 40 Hz and lose energy quickly between 40-80 Hz. The frequency at which the main lobe of the em-seq spectrum loses all power will be denoted as the cutoff frequency (COF; e.g., 80 Hz in the previous example). The frequency up to which the em-seq spectrum is approximately flat (i.e., loses only 2 dB in power) will be denoted as *f*_2*dB*_; thus, *f*_2_*dB* is 40 Hz in the previously given example. Given the properties of a sinc function, COF = 2\**f*_2*dB*_. A COF or *f*_2*dB*_ is set by choosing the elongation duration applied to the conventional m-seq to obtain the em-seq. In addition, the number of bits for the em-seq is a parameter that will need to be chosen, as is the case with a conventional m-seq. The minimum requirement is that the total length of the em-seq needs to be longer than the expected impulse response.

Two different em-seqs were used for characterizing brainstem (via ANF coincidence) and cortical responses (via EEG) given that the two systems were expected to reflect very different temporal coding abilities. The brainstem em-seq was a 9 bit m-seq with T =2 ms, so *f*_2*dB*_ was 250 Hz. A minority of units were measured with the elongation parameter set at 1 ms in duration, i.e., *f*_2*dB*_: 500 Hz; the data from these units indicated that *f*_2*dB*_ of 250 Hz was sufficient. The brainstem em-seq was 1.026 seconds in duration and was presented 6 times. The cortical em-seq was 8 bits with each point being elongated to 50 ms, giving a *f*_2*dB*_ of 10 Hz. The 50 ms duration for each em-seq point was chosen after piloting using *f*_2*dB*_ of 20 Hz suggested cortical phase locking limits to binaural modulations measured with EEG were below 10 Hz. The total duration of the cortical em-seq was 12.75 seconds, and we aimed to collect 300 trials for BMs applied to ITD and IAC. One participant did not complete all trials for the ITD em-seq, and one participant did not complete all trials for the IAC em-seq due to availability constraints.

Fig. 4.B depicts the paradigm used in this study. An em-seq modulated either the ITD or IAC of a broadband noise stimulus. The noise stimulus used for single-unit data had a bandwidth of 0.01 – 20 kHz and 0.2 – 1.5 kHz for the cortical stimulus. The bandwidth for the cortical stimulus was narrower since humans do not appear to use fine structure binaural cues for spatial perception beyond approximately 1.5 kHz (Brughera et al., 2013). An amplitude modulation (AM) was imposed at the highest frequency the em-seq can characterize (so 500 Hz for the brainstem em-mseq and 20 Hz for the cortical; matching the fastest rate at which the em-seq was allowed to switch states) to mask any phase discontinuities introduced by jumping between two ITD or IAC values. This does not affect our analysis because the em-seq would have lost most of its power at this frequency. The modulation by the em-mseq was binaural, so the em-mseq binaural modulation could only be heard with both earphones in place. Listening with just one earphone would lead to perceiving just 20-Hz amplitude modulated noise. The m-seq for IAC bounced between an IAC of 1 and −1, and for ITD bounced between 0 and 500 *μ*s for EEG data collection. For single-unit data, the ITD bounced between 0 and 1 / (2*CF), where CF is the characteristic frequency of the unit being measured. This was chosen based on the observation that ITD tuning curves of brainstem single neurons exhibit a trough at an ITD that is a phase of pi from the CF of the unit (Harper and McAlpine, 2004), and because we would expect greatest probability of coincidence between left and right auditory nerve responses for an ITD of 0, and minimum coincidence at a ITD corresponding to a phase difference of pi at CF.

**Figure 4.**
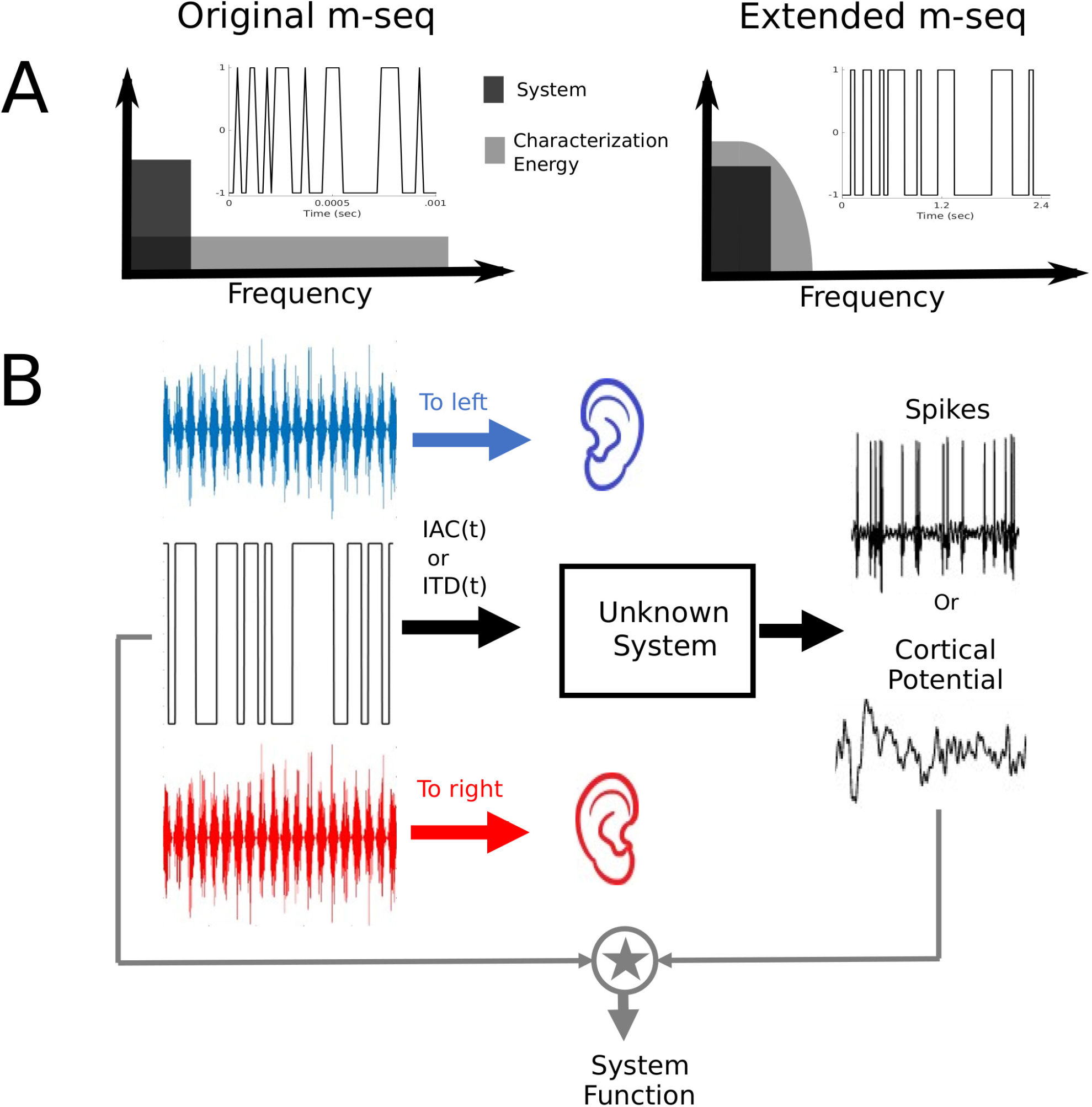
**A** depicts how the m-seq is transformed into the extended m-seq by increasing the duration of each point in the m-seq. This alters the frequency response of the m-seq to be sinc shaped instead of white, but that is useful in focusing the characterization energy in the range the system of interest is active. **B** depicts the paradigm used to obtain system responses. The extended m-seq modulated either the IAC or ITD of the noise stimulus and the neural measure, either spikes or voltage potentials (EEG), were cross correlated with the extended m-seq to obtain an estimate of the system response.

#### 3.4.1 Em-seq Analysis

The em-seq used for analysis will be termed the recovery em-seq (rem-seq). The rem-seq always goes between 1 and −1, so with IAC, the em-mseq played as the stimulus and the rem-mseq are the same. For ITD, however, the em-mseq bounces between two different ITD values, but for analysis the rem-mseq goes between 1 and −1. This choice of rem-mseq avoids introducing an artificial DC value into the analysis, and an overall scaling that would not affect the frequency shape of the system response, thus making it well-suited to characterize how the system can track modulations.

For the single-unit data, the spikes were treated as Dirac delta functions (i.e., single-sample impulses indicating that a spike occurred in a narrow time bin of the peristimulus time histogram) and cross-correlated with the rem-mseq to extract the binaural impulse response, i.e. binaural temporal response function. Noise floor estimates were constructed by randomizing the inter-spike intervals (ISIs) and making the first spike time be drawn from a unifrom distribution between 0 and the actual first spike time for that trial, and then doing the same cross correlation analysis on the jumbled spike train which has the same ISI distribution. See supplementary fig. 1 for examples of system functions obtained from single units utilizing this approach.

For the EEG data, the response in each channel was cross-correlated with the binaural rem-seq giving a multi-channel binaural temporal response function (mcBTRF) for each participant. The mcBTRF is then averaged across participants. In the mcBTRF, the response can be seen in several channels with different magnitudes and with different polarities. This is expected considering that the response from any given neural source will contribute voltage fluctuations in different EEG channels with different amplitudes and polarities depending on the geometrical configuration of the source relative to the sensors. To extract the dominant source and its spatial topgraphy, we used principal component analysis (PCA) on the mcBTRF. The PCA implementation from sci-kit learn in Python was utilized. PCA was done in the time range of 0 to 500 ms and only the first principal component was taken as the source binaural temporal response function, sBTRF. The IAC sBTRF explained 94% of the variance of the mcBTRF, and the ITD sBTRF explained 70% of the variance. The PCA operation on the mcBTRF can be thought of as a spatial filter that generates a linear combination of the 32 channels such that the combination (i.e., the sBTRF) accounts for most of the variance in the mcBTRF. The scalp topographic map of the PCA weights to get the sBTRF can provide information about physical location of the dominant source tracking the binaural modulations. To estimate the variance of the sBTRF, we used a jacknifing (leave-one-out) procedure. Noise floors for the EEG data were constructed by multiplying a randomly chosen half of the trials by −1 and then carrying out the same analysis as used to obtain the mcBTFRs. Ten noise-floor estimates were generated for each participant for each channel. The PCA weights from the actual mcBTRF were used on the noise-floor estimates to obtain the noise-floor estimates for the sBTRFs. See supplementary Fig. 2 for a visual depiction for going from the 32-channel evoked response to the sBTRF.

### 3.5 Psychoacoustic Experiments

We performed four psychoacoustic experiments, laid out in Fig. 1. All stimuli were generated using custom MATLAB scripts and delivered through the acoustic apparatus described previously. The FM tracking psychoacoustic experiment was done through our online platform that has previously been validated to yield average absolute thresholds that are comparable to lab-based data (Mok et al., 2021). For every task, participants were first given a brief demo to understand the task.

#### 3.5.1 Perceptual limits for detecting binaural modulations

To measure human ability to detect binaural modulations, we used the method of constant stimuli with the oscillating-correlation (OSCOR) stimulus. The OSCOR stimulus consists of noise tokens with sinusoidally varying IAC, and has been used previously in both behavioral and physiological studies (Grantham, 1982; Joris et al., 2006; Siveke et al., 2008). Each trial was 3-interval 3-alternatives-forced-choice with the target interval containing the OSCOR stimulus and the other two intervals containing interaurally uncorrelated noise (IAC =0). We evaluated performance at octave frequencies between 5-320 Hz with 20 trials at each frequency. The OSCOR stimulus was band-limited between 0.2 – 1.5 kHz because of data suggesting fine-structure-based binaural cues may not be useful beyond 1.5 kHz (Brughera et al., 2013). However, we repeated this experiment in one subject with white noise due to physiological data indicating cells with higher center frequencies can encode the fast OSCORs (Joris et al., 2006). Indeed, one possibility is that fine-structure-based binaural cues may be detected for higher (beyond 1.5 kHz) carriers but that these cues don’t inform spatial perception. The results of the measurement in the one participant with OSCOR applied to bandlimited (0.2-1.5 kHz) and to white noise (extending up to half the sampling rate) are shown in Supplementary Fig. 3.

#### 3.5.2 Limits for perceiving dynamic space

Several studies have anecdotally reported that with the OSCOR stimulus and other dynamic binaural stimuli, the perception of the stimulus appears to change from a spatialized image (i.e., moving in space) to a flutter around 6-10 Hz (Grantham and Wightman, 1978; Siveke et al., 2008; Zuk and Delgutte, 2017). We hypothesized that this switch would align with cortical temporal coding limits. Accordingly, we formally measured this switch in 1 participants using the method of limits with the OSCOR stimulus. There were 10 ascending and descending trials that started randomly between 3-6 Hz or 16-19 Hz respectively. The participant pushed a button indicating whether the perception of the stimulus had changed or not (either spatial to flutter or flutter to spatial) in each trial. If the perception had not changed, the frequency was increased (ascending trials) or decreased (descending trials) by 1 Hz until the change was noted.

#### 3.5.3 Perceptual dynamics of spatial unmasking & comparison to physiology

The third behavioral task probed dynamic binaural unmasking, and was based on a previously published paradigm (Culling and Summerfield, 1998). In this task the noise is uncorrelated (IAC =0) except for a window of time in the middle of the stimulus where the noise becomes completely correlated (IAC =1), see Fig.3 C. While the noise is completely correlated, an anti-correlated (IAC = −1) 850 Hz tone, 20 ms in duration, is played coincidentally with correlated noise. The difference in IAC between the tone and the noise (i.e., the “*N*0*Sπ*” configuration of the mixture) can be used to improve detection of the tone, i.e. a spatial unmasking effect. We varied the duration of the completely correlated period of the noise and measured detection thresholds for the tone using an adaptive 2-up-1-down paradigm. The window durations we evaluated were 0, 50, 75, 100, 125, 150, 200, 400, 800, and 1600 ms. Culling and Summerfield (1998) used this task to estimate what the underlying binaural temporal analysis window by comparing the unmasking function (dB masking release vs. window duration function) with levels of unmasking that different window shapes would predict. Here, we measured the binaural temporal window physiologically using EEG. Thus, instead of fitting arbitrary window shapes, we analyze how well the physiologically measured temporal window, the sBTRF, quantitatively explains the entire behaviorally measured unmasking function. This was done in two steps. First, the sBTRF (normalized and shifted to sum to 1 and take non-negative values) was convolved with the background noise, and the maximum “internal” IAC of the noise is estimated in the window of overlap with the tone. Then a binaural masking level difference (BMLD), or detection improvement relative to a window duration of 0 is estimated from the known relationship between static IAC and BMLD (van der Heijden and Trahiotis, 1997), which is is captured in Equation 1 below. van der Heijden and Trahiotis (1997) found that this equation could account for 98% of the variance of behavioral BMLD data from Robinson and Jeffress (1963). Here, *T*_*No*_ is the mean threshold at the largest window size (1600 ms) and *T*_*Nu*_ is the mean threshold with no window present.

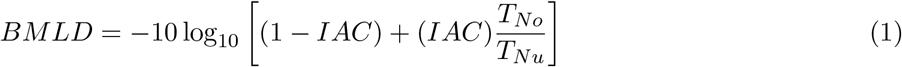

#### 3.5.4 FM phase difference detection using web-based psychoacoustics

In response to the COVID19 pandemic, we developed and validated a web-based platform for conducting suprathreshold psychoacoustics experiments (Mok et al., 2021). We recruited 14 participants from Prolific in the 18-55 year age range. Each participant passed a headphone-use screening test, and a screening for normal hearing based on a suprathreshold speech-in-babble paradigm (Mok et al., 2021) before participating in the main FM experiment. One of the authors also completed the task, yielding a total of 15 total participants.

In the main task, participants were instructed to detect the difference between two frequency modulations at a given modulation rate, but applied to spectrally distant carriers. One carrier was always between 500-750 Hz, and the other carrier was chosen to be two octaves higher than the first. The modulation depth of the FM was 10% of the carrier frequency. The FM rates we evaluated were 4,8,16,32, and 64 Hz and the phase difference between the FMs were 30, 60, 90, or 180 degrees. An example of the FM phase difference detection stimulus is shown in Fig. 4. The stimulus duration was 1.5 seconds and had a sampling rate of 44,100 Hz. To eliminate potential onset effects in detecting the phase difference between the two FMs a discrete prolate-spheroidal sequence (DPSS) window was used to apply a 125 ms ramp, and the starting phase of the FMs in each interval was randomized. Each trial was organized in a 3-interval 3-AFC format, with non-target stimulus intervals containing in-phase FMs and the target interval containing the FMs with a phase difference. Mean and standard error parameters for detection accuracy were estimated using the median, and the median absolute deviation, respectively.

---

**Supplementary Figure 1.**
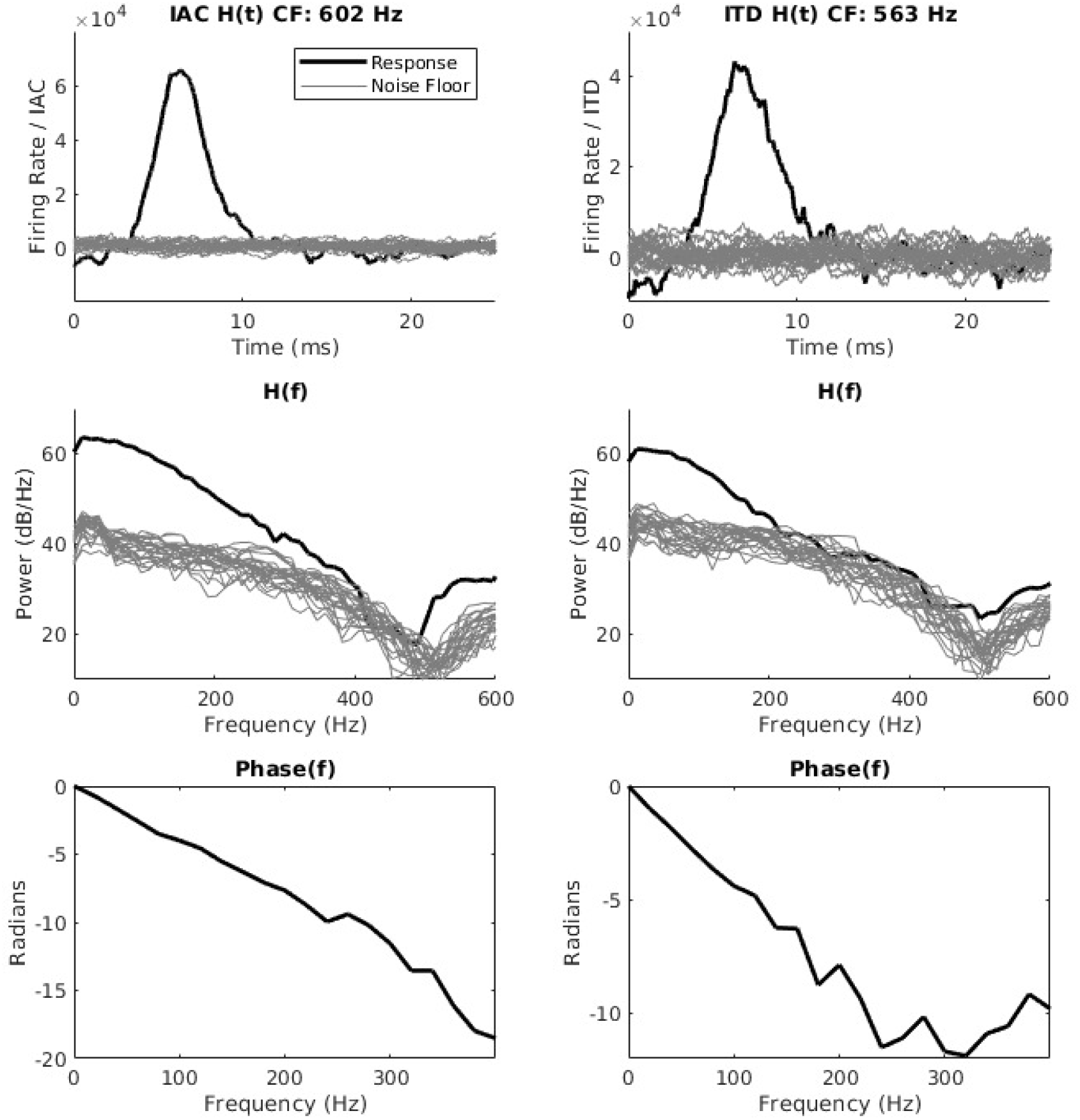
The left column contains an example impulse, frequency, and phase response from a unit with a center frequency (CF) of 602 Hz for IAC and the right column for ITD of a unit with a CF of 563 Hz.

**Supplementary Figure 2.**
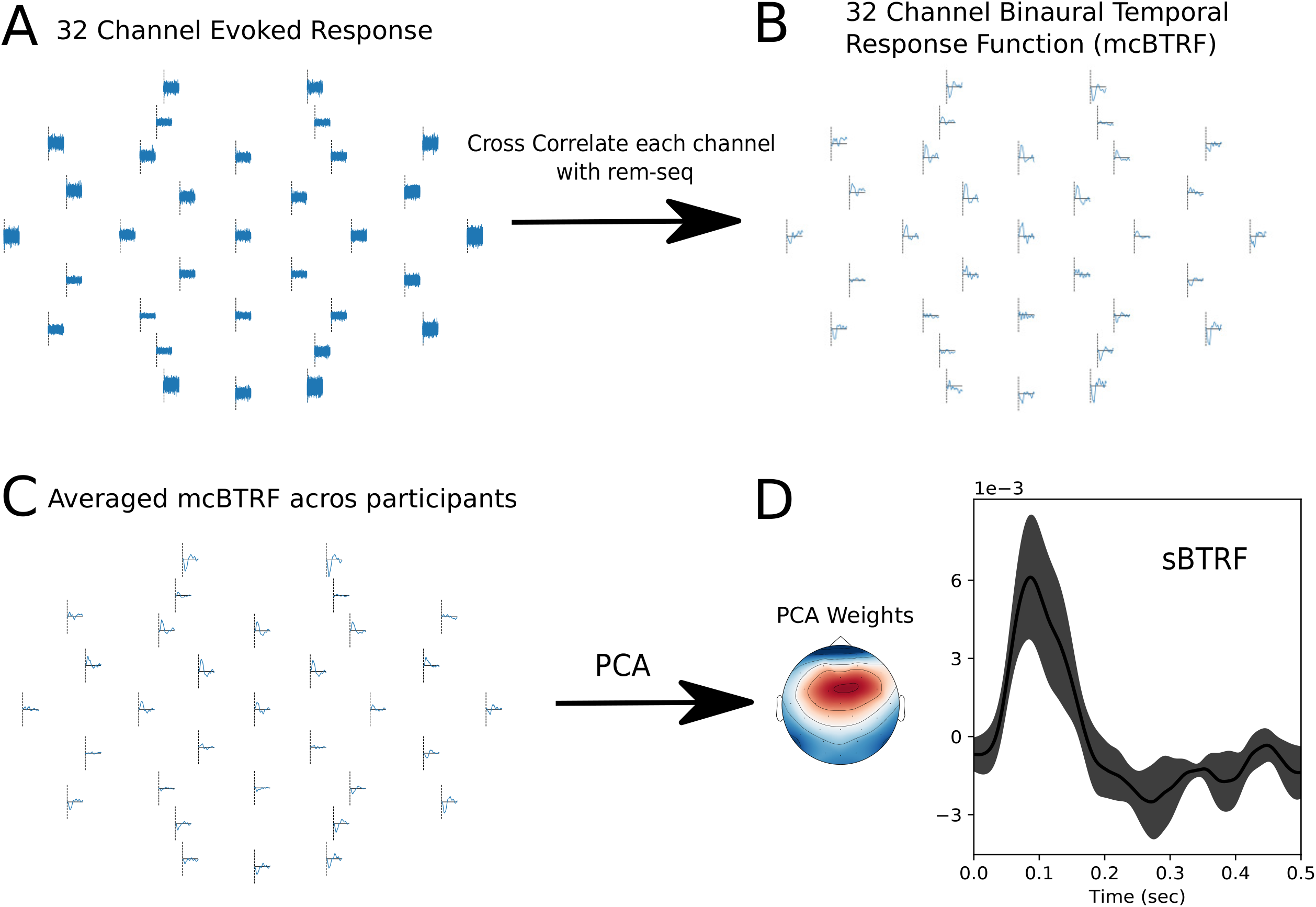
The figure shows the procedure of going from the 32 channel evoked response to the sBTRF. First each channel is cross-correlated with the rem-mseq to obtain the mcBTRF shown in **B**. The mcBTRF was averaged across all participants (**C**), and then PCA was done on averaged mcBTRF. The PCA weights and sBTRF which is the first principal component is depicted in **D**.

**Supplementary Figure 3.**
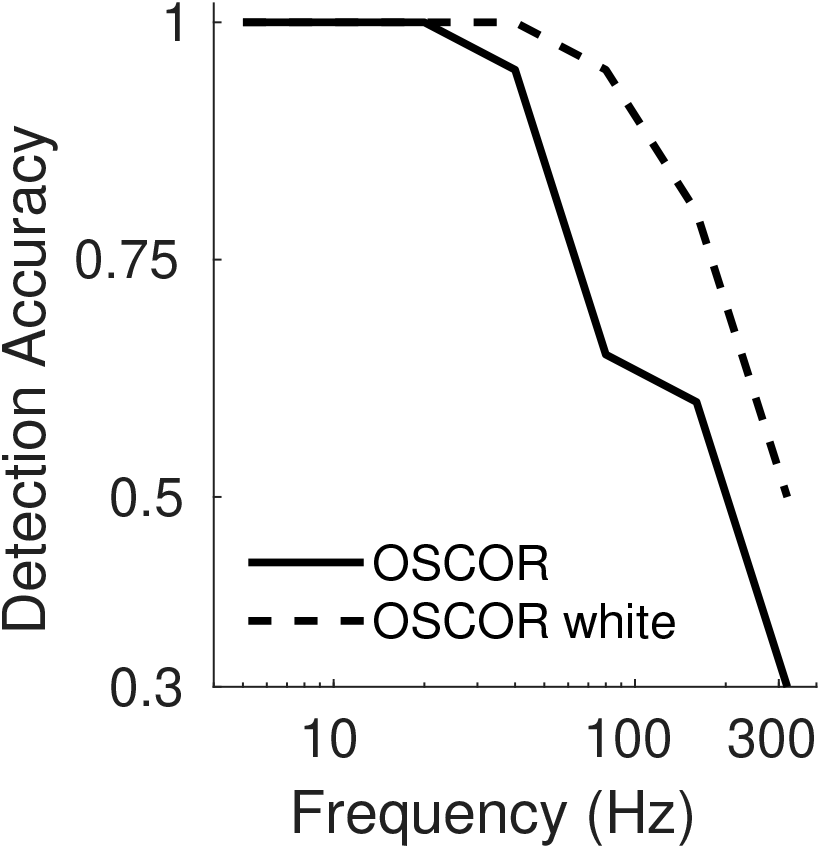
Results from one participant where the OSCOR was measured with the noise band-limited to 0.2-1.5 kHz and with white noise. The OSCOR can be detected out to much higher frequencies with white noise than band-limited noise.

**Supplementary Figure 4.**
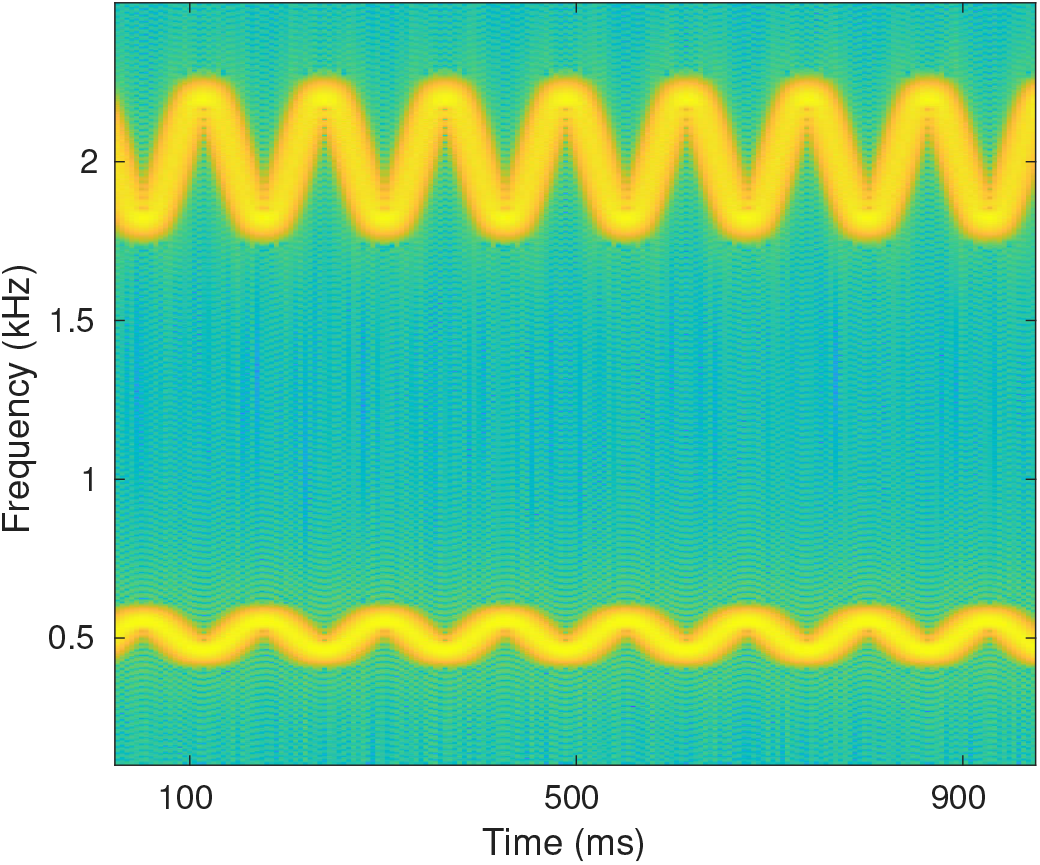
An example of the FM phase-difference-detection stimulus. In this example, the carriers are at 0.5 and 2 kHz, and the FM rate is 8 Hz with a phase difference of 180 degrees.

## 4 Acknowledgements

We would like to acknowledge Mark Sayles for his contributions to the development and execution of experiments in chinchillas to study brainstem level encoding of dynamic binaural cues. Mark Sayles also contributed to the development of the novel binaural systems identification approach used in this work. This work was supported by National Institutes of Health (NIH) grants R01DC015989 (HMB), and T32DC016853 (RS).

## Code and Data Availability Statement

The code to reproduce the figures in this work is publicly available on GitHub (https://github.com/Ravinderjit-S/DynamicBinauralProcessing) and archived using Zenodo (Singh, 2021). The EEG data is openly accessible and archived using Zenodo (Singh and Bharadwaj, 2021).

